# STORM-Net: Simple and Timely Optode Registration Method for Functional Near-Infrared Spectroscopy (fNIRS)

**DOI:** 10.1101/2020.12.29.424683

**Authors:** Yotam Erel, Sagi Jaffe-Dax, Yaara Yeshurun, Amit H. Bermano

## Abstract

We introduce a robust video-based method for estimating the positions of fNIRS optodes on the scalp. The method is fast, requires no special hardware, and is intuitive to use with developmental populations. Co-registration is a crucial step for reliable analysis of FNIRS data, yet it still remains an open problem when considering these populations. Existing methods pose motion constraints, require expert annotation, or are only applicable in laboratory conditions. Using novel computer-vision technologies, we implement a fully-automatic appearance-based method that estimates the registration parameters of a mounted cap to the scalp from a raw video of the subject. We validate our method on 10 adult subjects and demonstrate its usability with infants. We compare our method to the standard 3D digitizer, and to other photogrammetry based approaches. We show our method achieves comparable accuracy to current appearance-based methods, while being orders of magnitude faster. Our fast registration facilitates more spatially precise fNIRS analysis with developmental populations even in unconventional environments. The method is implemented as a open-source toolbox at https://github.com/yoterel/STORM-Net.

## 1 Introduction

Functional Near-Infrared Spectroscopy (fNIRS) is an emerging neuroimaging technique that is becoming the preferred method for acquisition of cortical activity in developmental and clinical populations [10, 17]. Its wide usage in developmental research is due to three main advantages. First, fNIRS provides better comfort for both the participants and the experimenter compared to fMRI. Second, it allows higher spatial separation compared to EEG. Third, it is less susceptible to participant’s motion during the study compared to both fMRI and EEG. These advantages make fNIRS the ideal method to measure the brain response under relatively ecological settings. Nevertheless, the level of confidence and acceptance of this neuroimaging technique is limited due to the lack of a standard method for inferring the underlying cortical region which provides the signal for a specific channel. There are two challenges in pursuing this standard method. First, the need for an exact estimation of the scalp placement of the probes. Second, the requirement for a standard age-dependent cortical template that matches each scalp placement with a specified underlying cortical structure (e.g., gyrus or Broadmann area). Previous studies have already estimated the probable underlying cortical structure for scalp locations [20, 12]. Recent studies have also established the age-dependent template that should be used [7] and even corrected for head size variations [25]. However, the first challenge has been relatively overlooked, or solved only by constraining the portability of the system and reducing its usability with developmental or clinical populations. These constraints include: the requirement for attentive motionless subjects [8], a metal-free environment [1, 16], a setup that constitute a contraption unfit for an infant [18], a need for specific illumination (mono-chromatic background, well lit area, strongly colored cap, etc.) [9], post-experiment registration [4] and intensive expert manual labor [14, 6].

In general, contemporary methods for estimating optode locations on the scalp are either hardware based or photogrammetry based. In the hardware based approaches, the most popular device types are electromagnetic localizers (e.g. [5]). These systems offer great accuracy but are limited by interference from metal objects around the participants, and require them to remain motionless for up to several minutes during the procedure. Early developmental and numerous clinical populations fail to meet this requirement. Moreover, the system itself is costly, and offers limited portability. Similarly, other hardware based solutions such as 3D scanners or geodesic arrays of cameras usually suffer from some or all of the same disadvantages (e.g. [18]).

Photogrammetry-based approaches use optical devices such as cameras to acquire (possibly multiple) 2D representations of the system, and register it relative to some known ground truth. Since they use the visual appearance of the cap as their source of information, their advantages include portability and simplicity of hardware requirements, and alleviating the main constraints posed by the 3D digitizer as described above. The method used by Lloyd-Fox et al. [14] is an example of using images to register the fNIRS cap, but they require expert manual annotators to mark positions of fiducials. Hu et al. [9] perform registration with the SfM algorithm for 3D reconstruction of a scene based on multiple images from different angles. However, their method still requires manually selecting the sensors’ location on the 3D model after reconstruction. Still, this offers much greater convenience and does not require any manual measurements to be performed. Furthermore, the algorithm heavily relies on good selection of matching feature vectors extracted from the set of images taken and on the SIFT algorithm to find them, which can yield inferior results under various lighting and filming conditions. This is especially noticeable for footage lacking a feature rich environment [26], such as the black cap used by Hu et al. [9]. Jaffe-Dax et al. [11] use a similar approach, but overcome the manual annotation with a template matching algorithm thus achieving a fully automatic solution. To counter the feature matching problem mentioned above, the authors printed a feature rich pattern on the cap. However, since the SfM algorithm takes significant time to complete, the fully automated system fails to reach its full potential: Infants may lose attention during this time, move abruptly, thus preventing the required images for the reconstruction to be successful. In addition, this method can only be used as a post-processing step as opposed to a possibly more desired on-the-fly configuration. For both methods described, the profile of image capturing required is extremely difficult to achieve with developmental population, since they require the participants’ focus to be away from the moving camera through most of its path. Drawing their attention to some other place while filming is challenging even for experienced and proficient neuroscientists.

Despite their convenience, state-of-the-art works still suffer from sensitivity to changing illumination conditions, relatively complex acquisition instructions, and most importantly, a lengthy and cumbersome registration process.

In this paper, we introduce a new registration method, alleviating said issues. In the heart of our solution lies STORM-Net, a Convolutional Neural Network which predicts cap orientation from raw video. Our registration process is performed in under 10 seconds, allows obtaining immediate results, requires nothing but a commercial low budget smartphone or camera, and is suitable for non-laboratory conditions due to its robustness. The method is validated through a series of experiments that measure accuracy, compared to the popular 3D digitizer approach[5].

## 2 Method

Our video-based approach is shown in Figure 1. Given a new cap, it is first registered in an offline step (i.e. without any human subjects present) for improved accuracy. This produces a network that is fine-tuned to the specific cap, and is utilized in real-time to register the sensors’ positions on a human subject. Taking a deep learning approach, we aim at a concise solution to the problem in an end-to-end manner. In other words, a single network that starts with a video and performs registration, all in one pass. This is desired when possible since it yields real-time performance and gives the network a holistic understanding of the problem (as opposed to exposing different networks to different steps in the process), thus is much less prone to errors. However, this approach poses a significant challenge: A suitably large data-set must be available since we need to estimate a much more sophisticated function over the input (which also implies larger training and prediction times). In our case, producing such data-set is not feasible at this scale. Realistically, one is able to create only a handful of videos, since their capture and annotation (i.e. through digitizer [5] measurements) are slow and tedious. For this reason, we chose a hybrid approach: our video processing pipeline is split into several blocks performing separate logical tasks. To decrease the amount of required data, some of the blocks use well-explored technologies that reduce the raw data (i.e. the video clip) into much smaller dimensions (2D “landmarks”) so annotation becomes significantly easier, and can be performed automatically. We still maintain the holistic view approach for the core network (STORM-Net) by exposing it solely to these landmarks which are theoretically sufficient to conclude the registration parameters using prior geometric knowledge of the system (Section 2.1.1). Furthermore, since using landmarks decouples the appearance and geometry of the cap (i.e., the whole image vs. 2D landmarks of particular 3D points) we can use synthetic data instead of actual videos, which dramatically improves the usability and adaptability of the method. Lastly, the modularity of the pipeline makes its predictions more explainable; if some prediction carries a large error, it is easier to detect which block failed and how to fix it compared to a one big black-box neural network solution.

**Figure 1:**
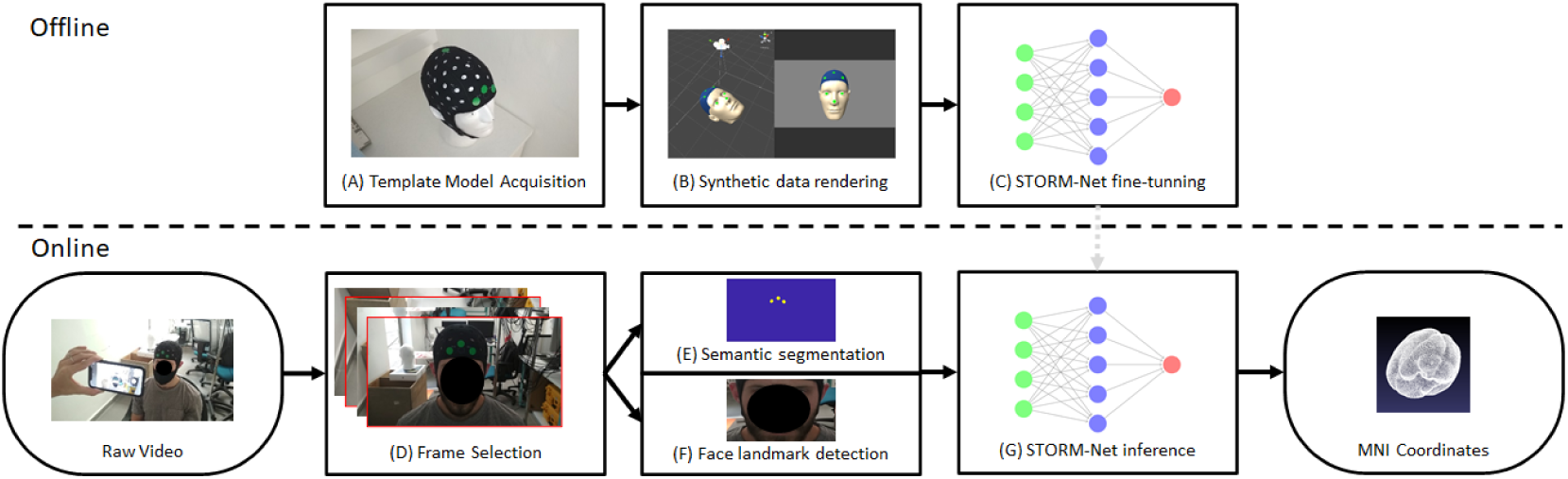
STORM-Net system overview. Given a new cap, the offline step is performed once for each cap configuration (top), while the online step is performed per subject in real-time (bottom). (A) Points of interest locations are acquired (e.g. by using a digitizer [5], or using the method introduced by Jaffe-Dax et al. [11]), (B) Synthetic data is automatically generated using a custom renderer (left: the virtual scene, right: projection of virtual scene to camera plane), (C) A STORM-Net model (tasked with estimating the cap-head registration using the three Euler angles) is trained on the synthetic data, (D) 10 frames are selected from a raw video, (E,F) Facial landmarks and cap fiducials are extracted from the video using a semantic-segmentation network and a face extraction model, (G) STORM-Net predicts the registration parameters and corrected locations of the points of interest can be retrieved (possibly also projected into MNI space if required).

In the offline step (Figure 1, top), points of interest locations (such as sensor or channel locations) are measured in 3D in a controlled environment using an external digitizer device [5], or any other 3D coordinate acquisition method (e.g. Jaffe-Dax et al. [11]) on a mannequin head (Figure 1, (A)). Then, a user-defined subset of these locations is used to produce synthetic 2D images, simulating a short video session (B). We use this data to train a rotation-estimation neural network (STORM-Net) that predicts the orientation of the entire cap from these 2D locations (C).

In the online step (Figure 1, bottom), the user takes a short video of a participant wearing the cap, from which registration is performed in real-time. The 2D locations of the aforementioned user-defined subset are then automatically extracted from the video (Figure 1, D,E,F), and are fed into STORM-Net (G). By applying the transformation predicted by STORM-Net to the template model, the final output is the corrected 3D locations of all points of interest, which can be projected to a desired coordinate system (such as MNI coordinates).

### 2.1 Offline Step

When using a new cap for the first time, the offline step is required. The user prepares an “ideal” environment where the cap is mounted on an anatomically correct mannequin head while being well aligned and positioned relative to it. The user then manually marks the positions of 4 or more specific locations (in all our experiments, we have used {“*Fp*1”, “*Fpz*”, “*Fp*2”, “*Cz*”}) on the cap by placing colored stickers on them. The 3D locations of these cap landmarks, 3 facial landmarks (the two eyes and the nose tip), along with any other points of interest on the cap (in all our experiments: all sensor locations), are then measured (e.g. by using an external digitizer device [5]). These 3D locations are referred to as the *template model* throughout the paper. The template model is fed into a renderer, which produces synthetic sequences of frames imitating the geometry and kinematics of a head wearing the cap in real videos. Each such sequence is a matrix of size 10 × 2*N*, representing a video with 10 frames where each frame has *N* (2D) positions of the cap and facial landmarks mentioned earlier (*N* = 7 in our case). The rendered sequence approximates the real video properties of the expected input in the online step. The produced samples are then used to train STORM-Net. Once trained, STORM-Net is able to predict the orientation of the cap relative to the scalp in all 3 axes in real-time. The entire training process takes approximately 30 minutes using a machine with a GPU available (or approximately 10 times longer without one). After performing the training step, only a negligible (less than 1 second) computational load is required for the online step (in addition to the handful of seconds required for the acquisition itself, as discussed in Section 2.2).

#### 2.1.1 Cap Preparation

As previously mentioned, we manually marked the positions of 4 cap landmarks, {“*Fp*1”, “*Fpz*”, “*Fp*2”, “*Cz*”}, using stickers. It is noteworthy that using these locations, or any standard anatomical landmark for that matter, is not necessary. Any sufficiently informative set of 3D positions is possible. Sufficiently informative in this case means that the matrix representing the set is at least of rank 6 (i.e. all 6 parameters of a rigid body in 3D can be deduced). The more landmarks used, the better the accuracy achieved (with diminishing effect), but measuring them makes the setup more cumbersome. Figure 3 visualizes the 3D to 2D landmark projection process. In our experiments, we used green colored stickers to mark the landmarks but this is not essential - their 2D positions are faithfully extracted by a semantic-segmentation network in the online step which was trained on merely 300 annotated photos, with very high reliability. A detailed description of the training process can be found in the supplementary material.

#### 2.1.2 Synthetic Data Generation

One of the biggest drawbacks of data-driven approaches in general, and neural network based learning in particular, is the need for a large body of example data to train on. We overcome this obstacle by introducing a framework that is based *solely* on synthetic data. To enable this, we train the network on landmark positions, instead of raw images, since those are easily imitated by synthetic means. Obviously, the prediction accuracy of STORM-Net on real videos is highly correlated with the ability to model and simulate them as best as possible. We use our renderer to perform this task, examples of which can be seen in Figure 2.

**Figure 2:**
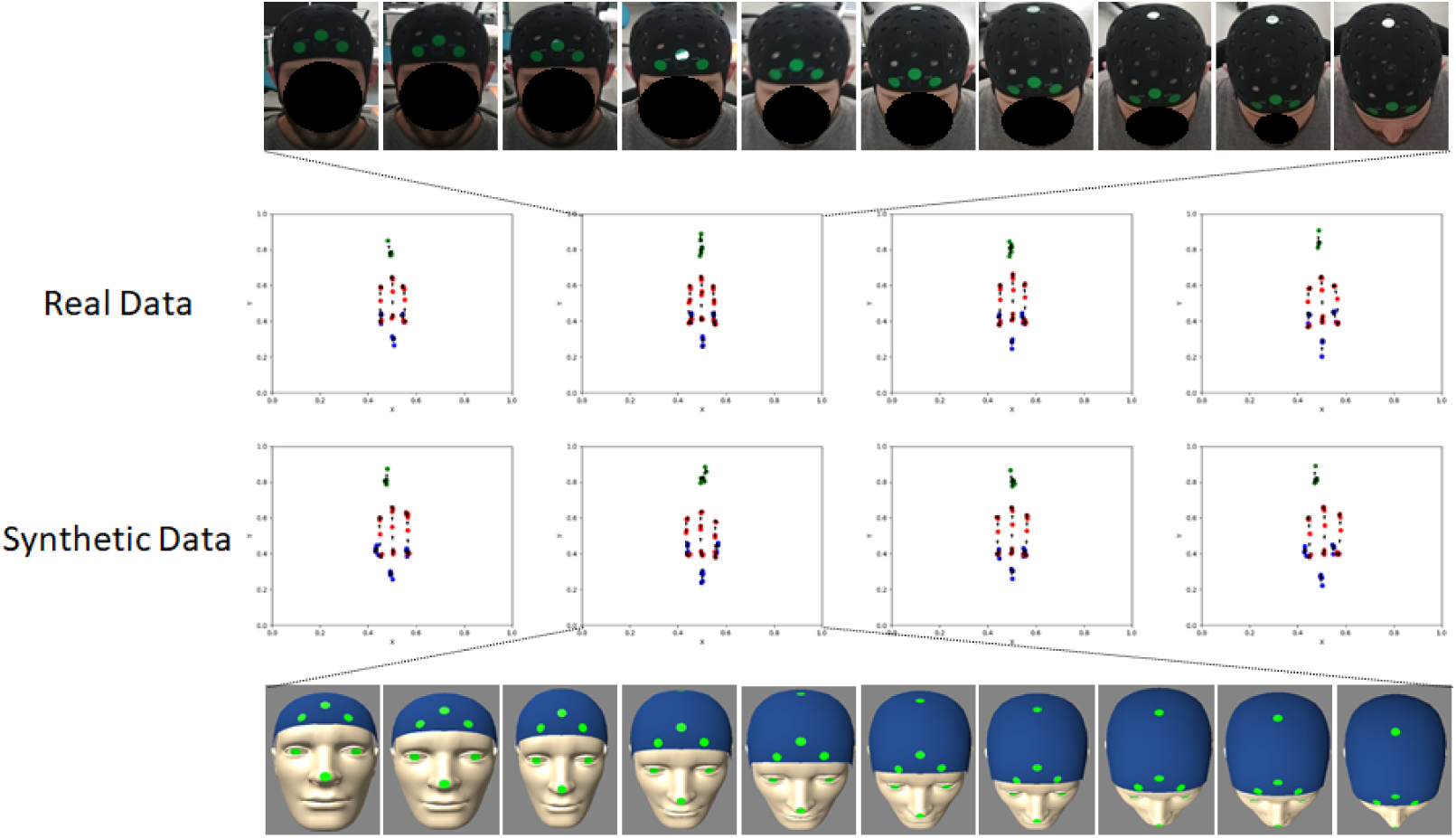
A visualization of real data (top) and similar synthetic data (bottom) extracted from different videos and rendering sessions respectively (each column). The points in the plots represent the 2D positions of the landmarks across a whole sequence. They are colored blue for facial landmarks, red for cap landmarks, and green shows explicitly the location of “Cz” for reference point. The data is centered and normalized. Notice how the synthetic data closely mimics the distribution of the real data and its noise sources. We used these graphs to aid with modeling the noise sources during scene creation in the renderer.

Using our renderer and the coordinates measured according the process described in Section 2.1.1, we generate sequences of 2D locations that emulate the capturing process described in Section 2.2. Since the human measuring process can never be ideally accurate, the most crucial part of the data synthesis process is correctly expressing the distribution of possible human input. The first deviation from an ideal scenario is that the camera never follows the exact required path, depicted in Figure 3. We handle this using 3 techniques. Firstly, we add random noise to the camera’s position and orientation during simulation (which we call the “Shaky Cam Effect”). Additionally, we add range deviations by increasing or decreasing the camera’s distance from the scene linearly with time. This is because we found that while taking the video it is often hard to keep the distance from the participant constant. Lastly, to mitigate the effects of the participants’ location being outside the center of the frame, we always center all landmarks using their center of mass as a post-rendering routine (i.e. after the data is rendered but before it is fed to the network).

**Figure 3:**
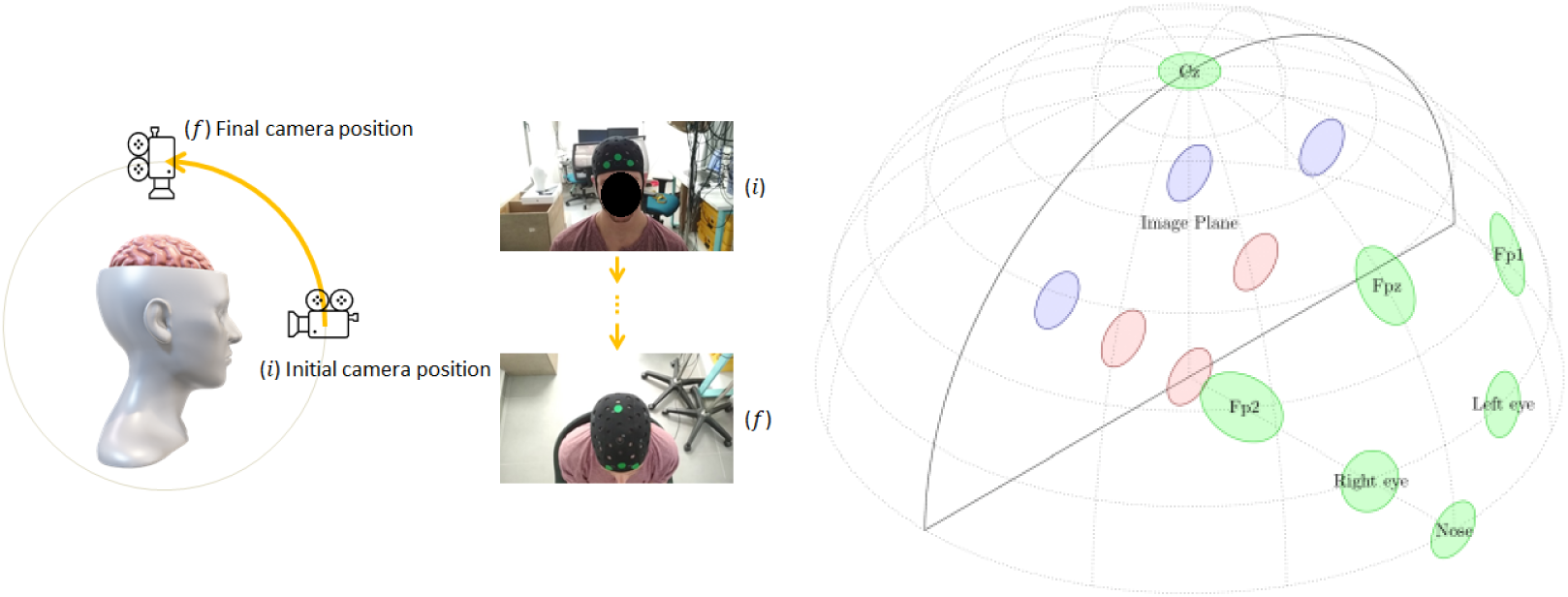
Left: The camera path designed for our system. The camera starts in front of the subject (i), and in a 5 second time frame it follows a circular path, reaching the top of the head (f). Note that the path is designed to be fast, efficient and resilient to subjects following the camera (since the camera moves beyond the head motion range). In practice even when infants followed the camera, the described path was easily achievable. Right: An illustration of projecting facial and cap landmarks to the camera’s image plane. Red projections indicate facial landmarks whereas blue projections indicate cap stickers (fiducials). Using a sequence of images taken from different directions, the data encodes sufficient information enabling the neural network to learn the orientation relative to scalp. Note that landmarks that are perpendicular to the image plane (such as “Cz” in the image) or landmarks occluded by the head are not visible in the projected image, causing loss of information that occurs during the capturing process, emphasizing the need for multiple images to be taken.

Another type of noise we take into account is partial occlusion. This happens because of the nature of the defined path, where the cap and face position relative to the camera occlude certain locations (e.g. the eyes are not visible when filming from above, which is the final position of the camera in the defined path). To handle this, we modeled a 3D representation of the face surface such that occlusions occur automatically by rendering the scene. We also artificially remove a random number of stickers from each frame of each data-point during training, inspired by popular drop-out techniques [24]. Hidden landmarks are placed at (*x, y*) = (0, 0), assuming the neural-network learns to ignore all such points given sufficient examples.

Lastly, selecting frames as discussed in 2.2.2 introduces temporal inconsistencies, such as non-smoothness, or even non-monotonic behavior, e.g., the cameraman momentarily shifts the camera in the opposite direction of the regular path. We handle such errors by randomly shuffling pairs of consecutive frames in the data-points.

### 2.2 Online Step

In this step, subjects are fitted with the same cap used in the offline step (with the stickers). The user then captures them on video using a camera (we used a low-budget smartphone [21]) that moves along a predetermined path for about 5 seconds. See Figure 3 for an exact description of the camera path. This video is then processed in the following way: we automatically select 10 frames from the video and feed them into an off-the-shelf facial landmark detector (based on a neural network) and our pre-trained semantic-segmentation neural network (which was trained to segment fiducials out of frames - green stickers in our case). These neural networks were designed to output 2D locations of the facial and cap landmarks mentioned above for every frame. Together their output consists of what is needed by STORM-Net to predict rotation parameters. We obtain these, and apply them to the template model to get the corrected positions of all points of interest. Optionally, we further project these into estimated MNI coordinates using the template model. As we demonstrate (Section 3), projection using the template model yields errors similar to those produced by the digitizer-based methodDigitizer [5].

#### 2.2.1 Video Capture

Our method uses raw video data to extract 3D information. Thus, it is essential this video includes useful data on the 3D relationships between the marked locations. Useful, in this case, means we are able to deduce the cap orientation and position relative to the scalp. As mentioned in (2.1.1), one way to make this possible is adding enough landmarks. Another useful way is using multiple images (video) of the same scene. To this end, we designed a camera path which follows a circular curve, ensuring these relationships are indeed captured when taking the video, while reducing interference with participant attention (which is imperative with the infant study group). Additionally, this path is easy to follow with a free hand held camera, without dedicated equipment, as demonstrated in Figure 3. In our case, videos were all taken using a low-budget smartphone [21] with 1920 ×1080 resolution and 30fps.

#### 2.2.2 Selecting the Frames

We found that using a somewhat naive selection generally suffices. We start by picking 10 frames evenly spaced from the video and naming them “base sites”. We observed that some specific frames are extremely blurry due to sudden movements of the hand held camera or the subject’s head. Additionally, the camera we used was of a rolling-shutter type and relatively of low performance. To mitigate this, we measured the amount of blurriness in a temporally local environment near every base site using the variance of the Laplacian operator [15], and selected the best candidate from every such environment (10 such selections are made). Nevertheless, this approach may fail if the video was taken based on a camera path significantly different than required.

#### 2.2.3 Extracting Facial Landmarks

The participant face is identified using an off-the-shelf facial landmark detector [13] as can be seen in Figure 4. We define a polygon around each eye’s landmarks, and take their average position as the eye’s “location”. For the tip of the nose, we use one specific landmark. If a face is not detected in some frame, then all coordinates of the facial landmarks are set to (*x, y*) = (0, 0) (for that frame).

**Figure 4:**
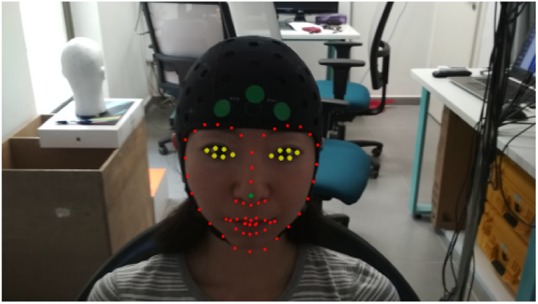
A frame with 68 facial landmarks extracted using an off-the-shelf extractor [13]. The eye landmarks (marked in yellow) are averaged to obtain the final eyes position and the tip of the nose (marked in green) is used as-is. Note that this landmark extractor is selected for its simplicity and effectiveness in our conditions, but can be seamlessly replaced by another.

#### 2.2.4 Extracting Cap Fiducials

Each frame selected is passed through a pre-trained semantic-segmentation neural network inspired by “capnet” [11] (see supplementary for architecture details and training parameters). The raw frame is then transformed to a binary mask where true valued pixels belong to the green stickers, marking specific cap locations. This binary mask is further processed into discrete 2D locations by locating their center of mass. To do this, we use the *findContours* method [2] to detect the blob contours in the binary mask and filter out blobs that are small (< 200 pixels in area for all our experiments), or located far from the center (>200 pixels away from the image horizontal center). These thresholds were determined empirically, but were observed to be robust in various scenarios. Finally, the center of the remaining four largest contours are reported as the cap landmark positions. If there are less than four remaining, the rest are set to *x, y* = [0, 0].

#### 2.2.5 Estimating the Rotation Parameters

In order to better exploit the convolutional power of neural networks, we first convert the landmark locations into heat-maps, a popular technique in the literature [3, 19], that encode the probability a landmark is located in any particular location. To do this, we treat the landmark locations as a matrix *F* × *N* where *F* is the number of frames (10 in our case), *N* is the number of landmark coordinates (7 · 2 = 14 in our case). For each frame, we define a 2D Gaussian mixture model distribution where the mean of every Gaussian is a 2D landmark location and the standard deviation is 5 pixels. We then sample this distribution over a 256 × 256 grid and obtain a heat-map. We stack all the heat-maps to obtain a 10 × 256 × 256 tensor, which is a single datapoint. We feed this data to STORM-Net, a 2D Convolutional Neural Network (CNN) that outputs 3 Euler angles {(*θ, ϕ, ξ*) ∈ ℝ^3^}. These are the rotation parameters that indicate how should the template model be rotated to fit the observed data. As mentioned in Section 2.1.2, STORM-Net is trained using synthetic data produced by our renderer (see supplementary for architecture details and training parameters). This synthetic data is also transformed into heat-maps using the same procedure. This representation encourages robustness to small mistakes in annotations and faster training rates compared to the more naive approach of using the matrix of landmarks directly.

### 2.3 Projecting into MNI Coordinates

The rotational parameters obtained from STORM-Net are used to rotate all the points of interest in the template model, which are then projected to MNI coordinates using the method of Singh et al. [20]. Note we use the anchors measured on the template model to allow the first stage of the algorithm to find an ideal affine transform to the common MNI template brain. This operation is an estimation of the real projection that needs to be performed using subject-specific anchors (at least 4 of them). All errors reported in this paper were calculated with respect to locations that were projected to MNI coordinates using this method (i.e. the results encompass the error that arises from this projection estimation). Additionally, to enable faster results, we implemented a new vectorized (parallel) version of the algorithm which runs 50 times faster than the current popular SPM for FNIRS toolbox implementation [23] using a GPU (or 10 times faster with a standard CPU). This solution, released along with the entire package, enables retrieving projection results in a matter of seconds even for a large amount of points of interest (90 sensor locations in our case).

### 2.4 Experiments

We conducted human experiments to validate our method. All experimental procedures were approved by the Tel-Aviv University Institutional Review Board for Human Subjects. All subjects provided written informed consent or parental informed consent for participating infants.

#### 2.4.1 Participants

10 subjects participated in this study (8 males, 2 females, aged 29.8*Y*± 2.73*Y* (*mean*±*STD*). Their head circumferences were 55.49*cm*± 2.14*cm* (*mean*±*STD*). Notice our selection of subjects with “medium” sized heads was made on purpose, so the cap we used fit well. This does not imply our method can’t work with other head sizes - this would simply require repeating the offline step as shown in Figure 5 for caps that better fit different head sizes.

**Figure 5:**
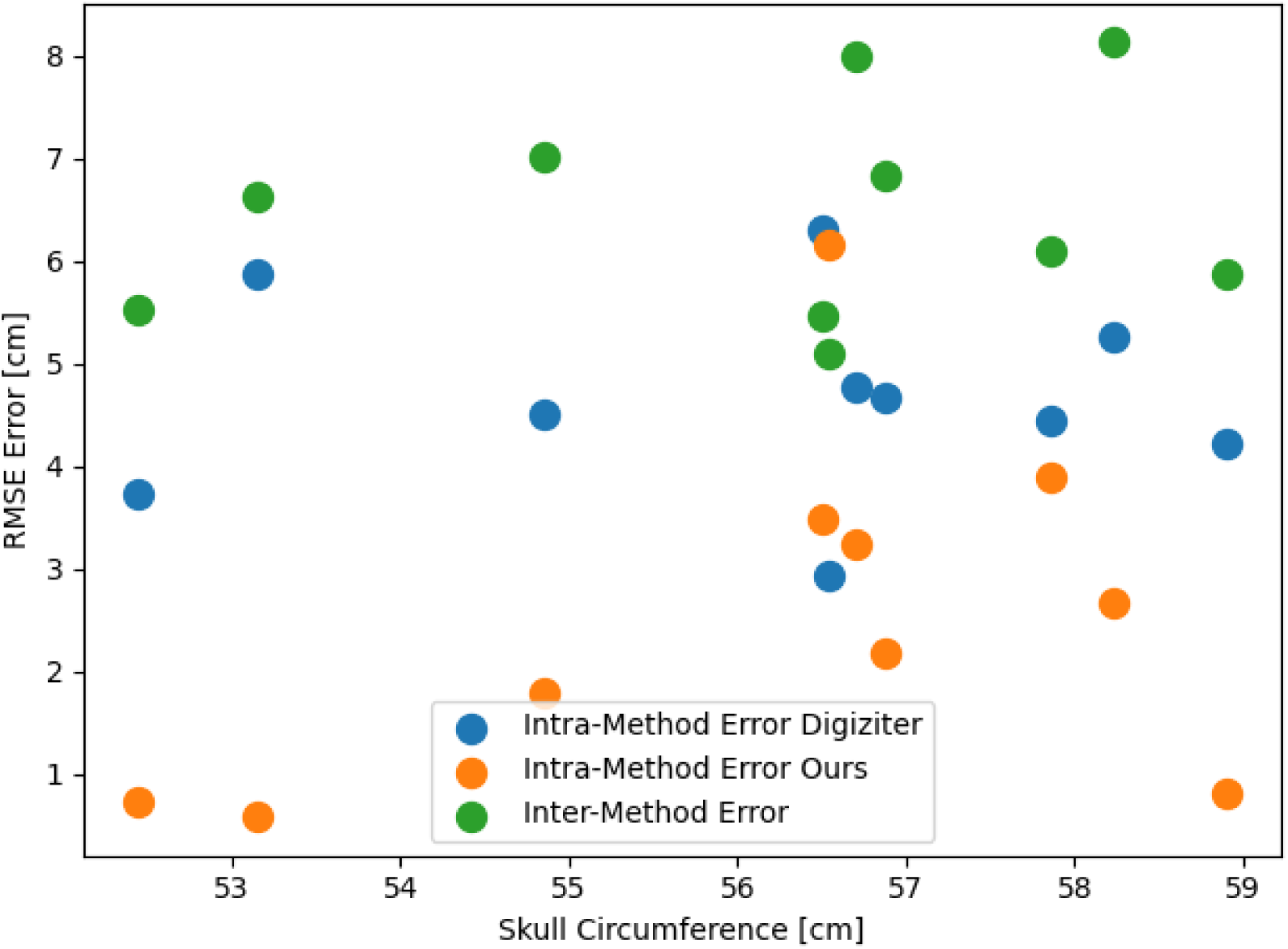
RMSE error as a function of skull size. The RMSE error for various skull circumferences of different subjects when using the digitizer twice (Intra-Method Digitzier), our method twice (Intra-Method Ours), and between the digitizer and our method (Inter-Method). No significant difference in errors can be observed across subject skull sizes using our method. This suggests that our method is robust to the size of the patients head size as long as the cap roughly fits.

**Figure 6:**
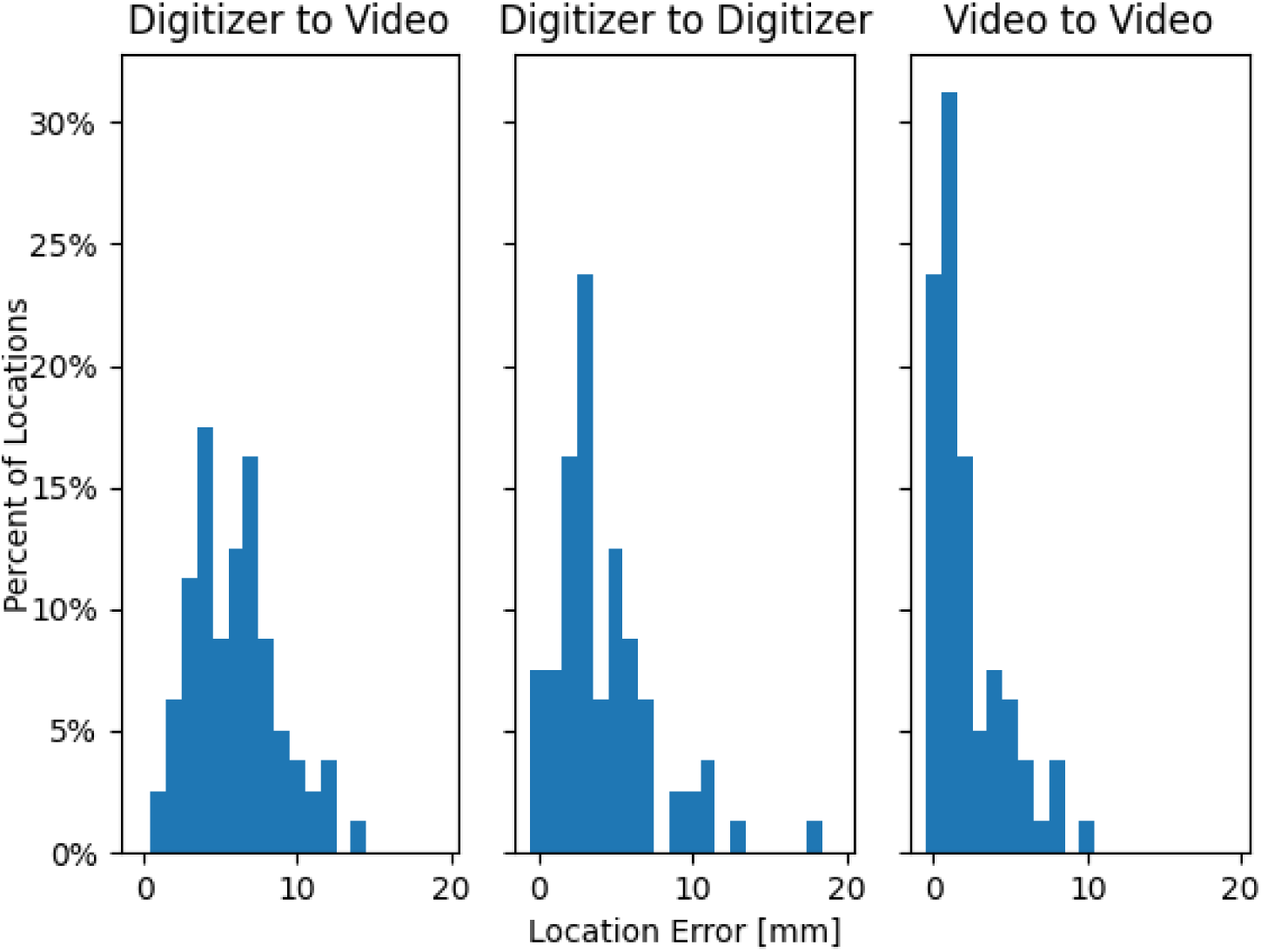
Distribution of errors for all measured locations. The errors were accumulated from all subjects using three comparisons. Left: 3D digitizer vs. Video (inter-method validation), Middle: 3D digitizer vs. 3D digitizer (intra-method reliability), Right: Video vs. Video (intra-method reliability). For a location visualization see supplementary material.

#### 2.4.2 Experimental Design

Participants were fitted with the cap which was well-positioned on their scalp (using standard positioning methods, without any external means). Two videos were taken followed by two digitizer measurements of certain landmarks while participants were wearing the cap - these were used to ascertain the intra-method reliability of the two methods and their inter-method validity. For the videos, this was done by feeding them to STORM-Net and comparing results. For the digitizer measurements, this was done using rigid body registration (Section 2.4.3). The cap was then removed from the scalp and re-positioned. Another video and another digitizer measurement for the landmarks were taken and were used to estimate the shift in positions of the channels by the two methods. It is worthy to note that digitizer measurements were done using 2 sensors, one fixed to the subjects’ head, and another used to do the actual measurements. This procedure ensured compensation for subjects moving their heads while the measurements are being done. However, it was observed to make little to no difference in the final results when using the second sensor (as subjects were relatively still). See supplementary material for more information.

#### 2.4.3 Rigid Body Registration

Given a rigid body model for the cap and two set of optode measurements *X*_1_, *X*_2_ ∈ *R*^*nx*3^; where *n* is the number of optodes and *X*_2_ was rotated and translated in some unknown manner, we use SVD decomposition [22] to obtain optimally aligned sets in the least squares sense. This was used to compute optode positions and shifts as measured by the digitizer for comparisons.

### 2.5 Implementation

Our software stack is written in python 3, and is freely available on Github: https://github.com/yoterel/STORM-Net

This makes it compatible with any operating system supporting python 3. The toolbox includes a GUI (graphical user interface) and a CLI (command line interface). The CLI allows a fully automated real-time end-to-end registration, while the GUI offers more subtle control over the process. The GUI includes a 3D model viewer (for visualizing the template model), a video annotation tool (users can manually or automatically perform annotation), the synthetic data renderer, and automatic training scripts for all described neural networks in this paper. Most importantly, it allows the registration of optodes locations in MNI coordinates from a given video file. Our implementation can process any video file format supported by the ffmpeg library which includes all popular file formats.

## 3 Results

### 3.1 Neural Network, Validated Through the Standard 3D Digitizer

Across all subjects (*n* = 10), measuring the same locations twice using our method yielded an intra-method reliability of 3.00*mm* ± 1.61*mm*; *mean* ± *STD* which was better than using the digitizer to measure those same location twice with an intra-method reliability of 4.67*mm* ± 0.93*mm*; *mean* ± *STD*. The distances between the positions of the same locations as estimated by the two methods were comparable yet slightly bigger to the distances between the positions of the same locations as measured by the digitizer twice (inter-method validity of 6.47*mm* ±1.32*mm*). However, the error of our method is overall still an order of magnitude smaller than the typical distance between optodes (3*cm*). All the subjects were fit with the same cap, and we have not observed any bias in prediction accuracy with regards to certain head sizes as shown in Figure 5.

### 3.2 Neural Network Cap Shift Detection

After placing the cap and performing validation measurements twice, as described in the previous section, the cap was removed and re-positioned on the subjects head, and new measurements were performed (Using a digitizer and our method). As expected, this re-positioning process introduced some shift in optode positions. We found a high correlation between the measured shift in optode positions using the digitizer and the measured shift using our method. Since we use a rigid model registration it is sufficient to describe such shifts using the rotational parameters *θ, ϕ, ξ* which represent the rotation around the X axis, Y axis and Z axis respectively. We obtained Pearson’s coefficients of 0.500, 0.381, 0.492 for *θ, ϕ, ξ* with *P* = 0.171, 0.312, 0.179, respectively, across all subjects, which indicate a positive close to linear relationship between shifts detected by the digitizer and shifts detected by our method.

### 3.3 Automatic Channel Estimation in Infants

We performed a qualitative evaluation of our method on infants using a different template model, and observed results which were consistent with our speculations, even for very large rotations of the cap relative to the scalp as shown in Figure 7. When the cap is positioned with an angle relative to the scalp, STORM-Net still easily predicts the rotational parameters for each video. These parameters can be used to register the cap and obtain the 3D positions of all optodes using a template model of the cap.

**Figure 7:**
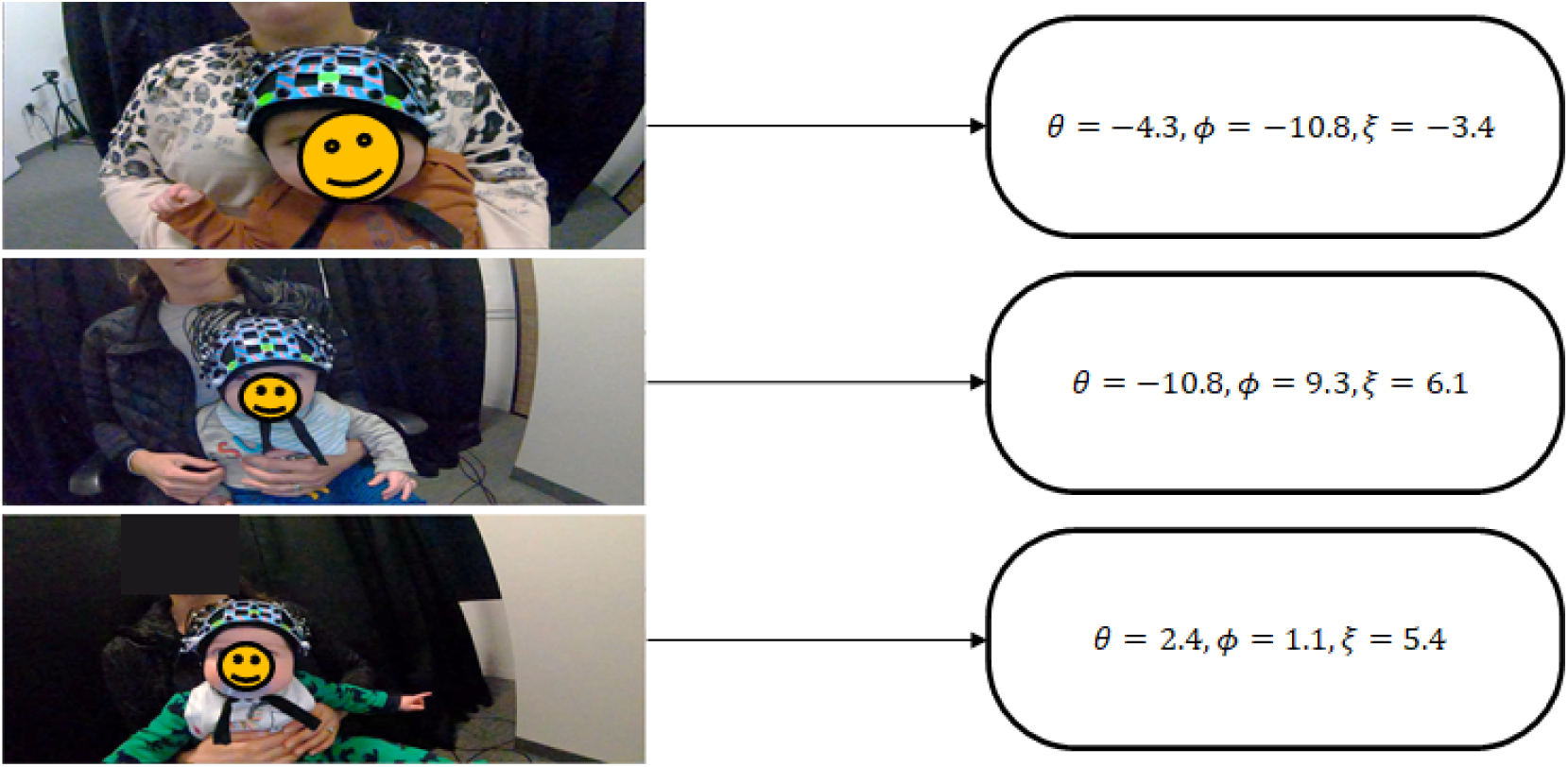
Qualitative evaluations of STORM on infants: The figure shows the 3 rotational parameters obtained using STORM from various infants videos. We observed that the parameters conform to our speculations in every axis across all videos we tested. For example, the first 2 videos (top 2 images) got negative and positive values for *ϕ*, which is expected for rotations around the *Y* axis (back to front). Notice the parameters are given in a right-handed coordinate system (i.e. angles can be interpreted as rotations around their respective axes, using the right hand rule).

### 3.4 Comparison to Previous Works

We compared our method to other photogrammetry based approaches used for registration of the optodes presented by Jaffe-Dax et al. [11] and Hu et al. [9]. Comparison was done based on the reported accuracy and speed of the method, the level of human involvement in the process of registration, and special equipment or limitations required by the method. Overall, our method performs well in all categories, as shown in Table 1. The acquisition and processing times of Hu et al. [9] were estimated based on the fact that they utilized the same algorithm as Jaffe-Dax et al. [11], adding an extra step of manually selecting the optodes locations on the 3D reconstruction. For the inter-method error of Hu et al. [9] we calculated the STD from their provided table. Hu et al. [9] use the SfM algorithm which relies on SIFT to perform feature matching, yet it does not perform well with visually uniform patches [26] and is, therefore, highly susceptible to the cap model. We assume the authors had to hand-tailor the visual appearance of the cap to a satisfactory degree, an action that requires some computer-vision expertise. Moreover, this would not be robust for other cap types or models. To overcome this challenge, Jaffe-Dax et al. [11] use visual Perlin noise patterns printed on the cap allowing the SfM algorithm to achieve good reconstruction. However, adjusting the cap properties still does not address other appearance based noise sources (e.g. occlusions, lighting conditions and quality of video) which affect SIFT greatly. Also this method forces stickers to be placed on the face of the subject, which is very distracting for the developmental study group. As opposed to these methods, we avoid the need to find good matching features all-together, while still providing robustness to visual appearances of the cap and the scene. This is done by using convolutional neural networks to semantically segmenting the frames, which are robust against appearance conditions because of their data-driven nature (the training process included augmenting the brightness of the scene). Thus, our method is useful for any cap in numerous different environments, as long as green stickers appear in the video and without the need to place them on the face of the subject. Retraining also makes it possible to work with other colors, and yield even better results if needed, by annotating as little as 300 images manually (the training procedure is described in the supplementary material). Finally, in contrast to the other solutions, our method’s speed allows fast re-registration on the spot when the taken video fails to meet the essential requirements (e.g. wrong camera path, the subject moved too much, occlusions, etc.). For adults, the cap can even be adjusted in real-time for better positioning on the scalp, using the registration results rather than using them as a post-processing step (for developmental population, once the cap is placed it is rather challenging to re-position it without distracting the subject).

**Table 1:** Comparison of photogrammetry based approaches

|  | Hu et al. [9] | Jaffe-Dax et al. [11] | **Ours** |
| --- | --- | --- | --- |
| **Intra-method error (adults, $AVG \pm STD$)** | Not Reported | $2mm \pm 0.5mm$ | $3mm \pm 1.6mm$ |
| **Inter-method error (adults, $AVG \pm STD$)** | $6.66mm \pm 3.30mm$ | $3.40mm \pm 0.90mm$ | $6.47mm \pm 1.32mm$ |
| **Acquisition Time**<br>**Processing Time** | $\sim 1$ minute<br>$> 10$ minutes | $\sim 1$ minute<br>$\sim 10$ minutes | $\sim 5$ seconds<br>$< 1$ seconds |
| **Automation of method** | semi-automatic | fully-automatic | fully-automatic |
| **Camera specifications** | 8MP resolution per image | 2.1MP resolution per image, 240 fps | 2.1MP resolution per image 30 fps. |
| **Environmental constraints** | mono-colored walls or cloth/curtains covering the background. | None | None |
| **Sensitivity to cap appearance** | High | Medium | Low |

## 4 Discussion

We proposed a real-time, automatic, photogrammetry-based method to register positions of optodes to the scalp. When positioning the cap on a subject’s head, even minute shifts in its orientation relative to the scalp might translate to large errors in the cortical regions being recorded. Such errors, highly common especially for developmental populations, propagate into the aggregation of signals between subjects as noise, and harm the process of conclusion drawing regarding cortical activity.

Our approach uses a pipeline consisting of state-of-the-art neural networks to correct these errors by registering the position of the cap relative to the scalp in an accurate, timely, and convenient manner. We designed this method to overcome many of the disadvantages of traditional techniques, effectively facilitating its usage for developmental populations. Our tool opens new possibilities in the field of cognitive neuroscience by allowing out-of-lab registration, and thus experiments to be performed with very little constraints. This is due to the nature of the data-driven convolutional neural-networks we use and their robustness to many scenarios. In addition, the method is automatic and performs in real-time, which means the user can get immediate feedback regarding the quality of the registration and easily perform it again if required.

We presented evidence that support the proposed method’s reliability and validity using the widely accepted field’s standard - the 3D digitizer. We also showed it is on par with previous photogrammetry based approaches in terms of accuracy, yet orders of magnitude faster, which aligns well with young children’s and infant’s tendency to lose attention easily.

We note that we have used a modified EEG cap for our validation experiments. Since EEG locations coincided with some of the optode locations, this means there is nothing to prevent this method from being used to register EEG caps to the scalp, and the errors presented hold for this scenario too.

To summarize, our easy-to-utilize registration approach strengthens the major advantage of fNIRS as being the optimal neuroimaging method to measure cortical response in naturalistic settings and developmental populations, by providing a real-time, automatic, and very simple to use method to register the cap to the subject’s scalp.

### 4.1 Limitations

We have modeled the transformation undergone by the points of interest after placing the cap on the scalp using a rigid-body model. In reality, this transformation is not guaranteed to be as such due to deformations and stretching undergone by the cap. To address this, we described an offline step which takes custom template models as input and fine-tunes STORM-Net to accommodate for a different type and size of the cap. This minimizes errors caused by cap deformations during inference, keeping the rigid-body model assumptions more faithfully. In addition, most fNIRS caps enforce a fixed distance between all optodes, which further constrains the type of possible deformations it can undergo. Thus, for these types of caps we expect even better performance than measured. Analysis of the spatial distribution of the errors across the surface of the cortex indicate a tendency for larger errors near the back side (See supplementary). We believe this can be mitigated by choosing more sticker locations in the back of the head. We also make an implicit assumption about the template model (specifically, the locations presumed to be anatomically correct) being a good proxy for the real subject-specific variability in scalp surface structure. This assumption is valid for a limited range of skull shapes and may break for skulls significantly different than the template model. However, we show this source of error across subjects is low enough to assure quality registration. Lastly, using neural networks as an estimation tool potentially entails unclear behavior, since we can never be sure how the decision for any particular registration was made, and how well it performs on a particular instance of the input. This is a well known and common problem to all neural networks. To address this, we split the task at hand (i.e. from raw video to registration parameters) into smaller discrete blocks that offer better explainability. This way, if the registration process fails for some reason, it is much easier to trace back what went wrong (and possibly perform manual corrections using the GUI) in this kind of design rather than using a black-box end-to-end neural network.

## Acknowledgments and Disclosure of Funding

All authors declare that they have no conflicts of interests.

This research was supported by the Israel Science Foundation grant 2434/19 to YY and Alon Scholarship to SJ. This research was also supported in part by Len Blavatnik and the Blavatnik family foundation to AB.

