## Supplemental Material for "STORM-Net: Simple and Timely Optode Registration Method for Functional Near-Infrared Spectroscopy (fNIRS)"

### STORM-Net - Supplementary

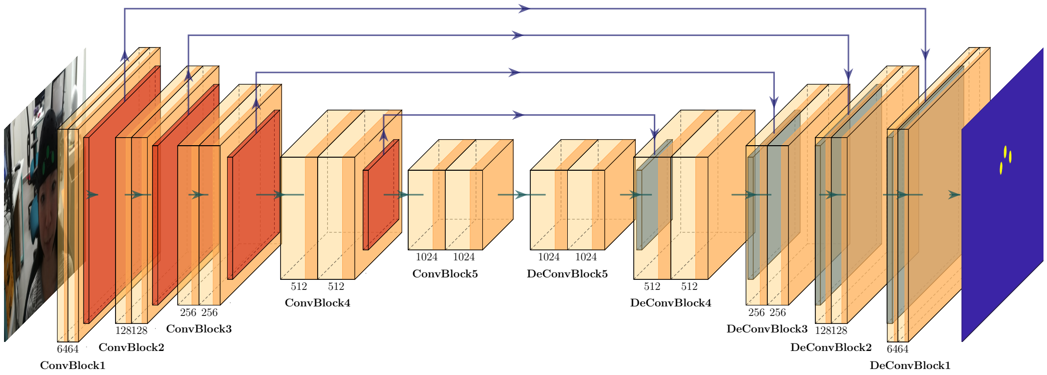


Figure 1: Our semantic-segmentation network architecture: The input is a tensor representing a video-frame (or a patch from it). The ConvBlocks in the image represent a 2D convolutional layer (with the number of filters indicated below, and a kernel of size (3,3)), which include a dropout layer (with dropout=0.2) between the 2D conv. layers, and a batch normalization layer after every second convolutional one. The down-sampling operations are performed using a MaxPooling2D layer (that pools using a factor of 2 in each axis), while the up-sampling operations are implemented using a Conv2DTranspose layer (with kernel size of (2,2) and stride size of (2,2)). Notice the skip connections between the encoder and decoder - these are typical for U-net–like architectures [2], and help achieve better results. We used ReLu activations in all cases except for the output layer, which uses a Sigmoid activation.

**Semantic-segmentation Network Detailed Description** Our semantic segmentation network is based on the popular U-net architecture [2] and inspired by "capnet" described in [1]. A detailed overview can be seen in Figure 1. Data was manually annotated for 300 random frames, a process which can be repeated for different colored stickers. We used a batch size of 8, and the Adam optimizer with a learning rate of $1e^{-5}$. Training was stopped once validation loss did not decrease for 20 consecutive epochs. Instead of feeding the whole images to the network, we split them into evenly spaced, non-overlapping patches of size 128x128 pixels. This improved network performance substantially, as the stickers on the cap amount to a negligible amount of pixels out of the whole frame, thus there is a great class imbalance in the data. We also perform various augmentations to the patches in the form of random rotations, flips, shifts, shearing and zooming in and out.

**STORM-Net Detailed Description** An overview of the architecture can be seen in Figure 2. We trained STORM-Net using batch sizes of 16, with an Adam optimizer with an initial learning rate of $1e^{-3}$ decreasing by a factor of 0.2 every 3 consecutive epochs with no improvement of the validation loss. Training was stopped after 100 epochs. The network architecture was: input->Conv2D(64)->BatchNorm->ReLU->MaxPooling2D->Conv2D(128)->BatchNorm->ReLU->MaxPooling2D->Conv2D(256)->BatchNorm->ReLU->MaxPooling2D->Conv2D(512)->BatchNorm->ReLU->MaxPooling2D->Flatten->FC(16)->ReLU->FC(3)->output, where the input is the (batch x 10 x 256 x 256) tensor of heat-maps, Conv2D(x) is a 2D convolutional layer with x output filters, kernel of size (3,3), and stride and padding of size 1, MaxPooling2D is a

Preprint. Under review.

down-sampling operation over the window of size 2 and stride 2, Flatten unfolds the inputs into a row vector and FC(x) is a fully connected layer with x outputs. Note the loss was defined as the mean squared error, since we regress to 3 continuous Euler angels (the output layer is a row vector of size 3).

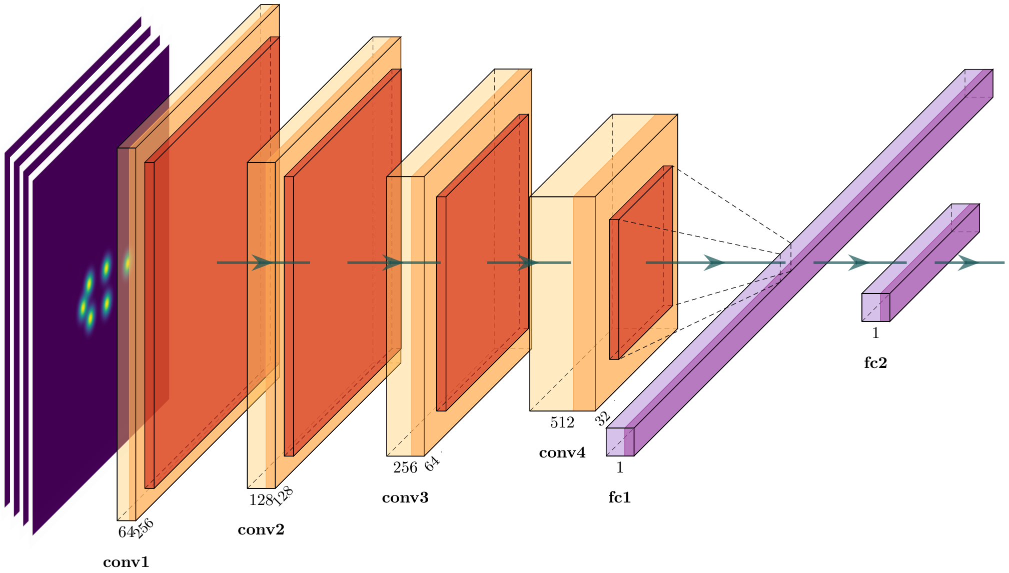


Figure 2: A schematic image of Storm-net architecture: The input is a stack of 10 heat-maps representing 10 frames with pixel values indicating the probability that the position is occupied by a landmark. The convolutional layers all use a kernel size of (3,3) with stride and padding of 1, with Batch Normalization layers after each of them. ReLU activations are used for all layers except the output.

**Robustness Test** We studied the sensitivity of the network predictions for randomly noised landmark inputs. The purpose of this was to ensure that similar user or automatic labeling of the landmarks yields similar results. Results are presented in Figure 3. In spatial terms, the results indicate that approximately for every $1mm$ error in the input, $1mm$ error in the output is induced.

**Digitizer Two Sensors vs. One Sensor** We verified that using the two sensors of the digitizer together was justified. These sensors include the pen-like sensor which allows exact localization by pressing a button attached to it and a second, static sensor which is attached to subject heads to allow compensation for their head movements during the experiment. Across all experiments, intra-method reliability when using the second sensor was $4.67mm \pm 0.93mm$; $mean \pm STD$, compared to not using the 2nd sensor yielding $4.94mm \pm 1.42mm$; $mean \pm STD$. Although using a second sensor improved accuracy, these differences are negligible, indicating that errors in digitizer measurements associated with head movement of the subject are rather insignificant compared to human errors associated with pinpointing an exact required location.

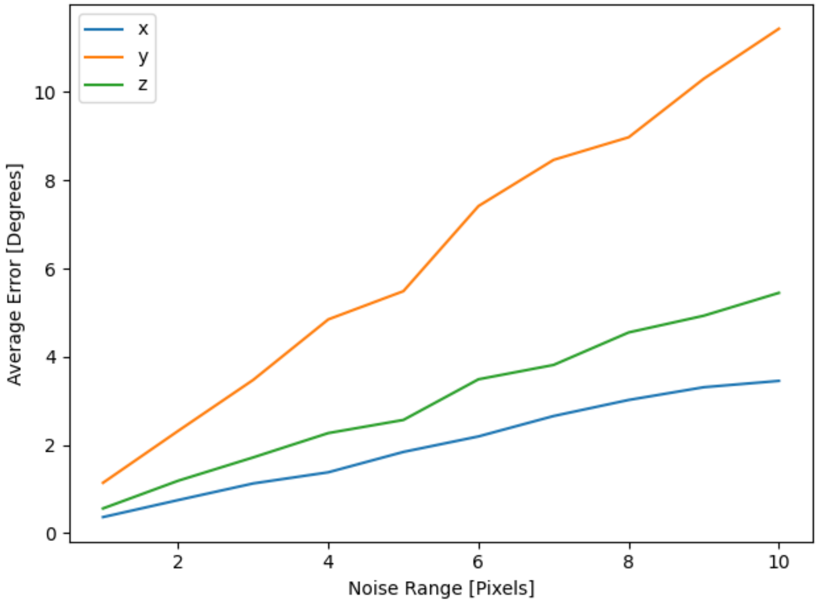


Figure 3: System robustness experiments: average error (in degrees) as a function of the noise added to the input landmark positions (in pixels). We add noise from a uniform distribution to each of the landmarks (noise is added per landmark from within the range $[-x/2, +x/2]$ where $x$ is the horizontal axis parameter). This was done 10 times for each $x$ value. The noised landmarks were inserted through STORM-Net to obtain rotational parameters, and the average error was measured. The network exhibits a close-to-linear error response to errors in the landmarks location. Interestingly, there is significant difference between the amount of error induced by each axis of rotation, suggesting STORM-Net is more sensitive to rotations around the Y axis (back to front).

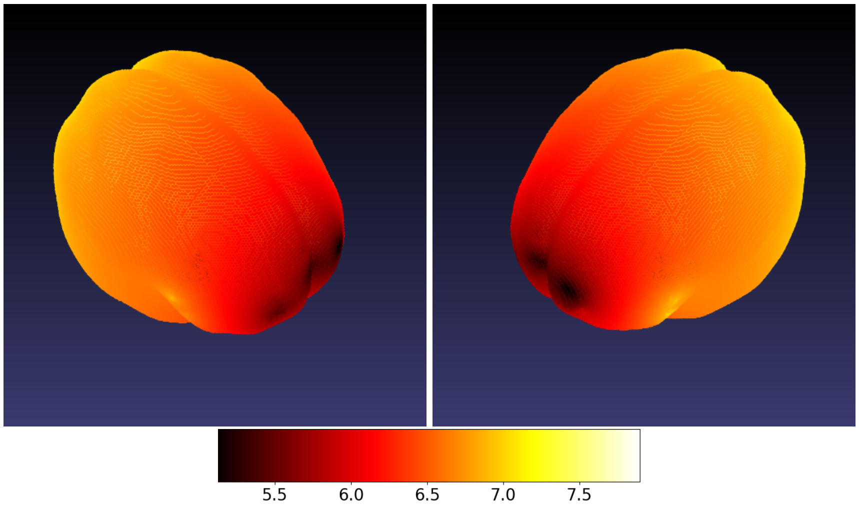


(a) Mean Inter-Method Validation Error [mm]

Figure 4: Spatial distribution of our inter-method validity errors displayed on an average MNI template brain. Errors in locations that weren't directly measured by the digitizer (i.e. for which no ground truth exists) were interpolated using a weighted average of measured locations. The average error across all subjects using this interpolation is $6.29mm \pm 0.27mm$; $mean \pm STD$. There is tendancy for higher accuracy near the frontal part of the brain. This is probably due to the abundance of visual cues (green stickers) present there which STORM-Net relies on in the offline and online steps.
